# Ndel1 promotes keratin assembly and desmosome stability

**DOI:** 10.1101/2020.04.30.070086

**Authors:** Yong-Bae Kim, Daniel Hlavaty, Jeff Maycock, Terry Lechler

## Abstract

Keratin intermediate filaments form dynamic polymer networks that organize in specific ways dependent on the cell type, the stage of the cell cycle, and the state of the cell. In differentiated cells of the epidermis, they are organized by desmosomes, cell-cell adhesion complexes which provide essential mechanical integrity to this tissue. Despite this, we know little about how keratin organization is controlled and whether desmosomes actively promote keratin assembly in addition to binding pre-assembled filaments. We recently discovered that Ndel1 is a desmosome-associated protein in differentiated epidermis. Here, we show that Ndel1 binds directly to keratin subunits through a motif conserved in all intermediate filament proteins. Further, Ndel1 is necessary for robust desmosome-keratin association and sufficient to reorganize keratins to distinct cellular sites. Lis1, a Ndel1 binding protein, is required for desmosomal localization of Ndel1, but not for its effects on keratin filaments. Finally, we use mouse genetics to demonstrate that loss of Ndel1 results in desmosome defects in the epidermis. Our data thus identify Ndel1 as a desmosome-associated protein which promotes local assembly/organization of keratin filaments and is essential for both robust cell adhesion and epidermal barrier function.

## INTRODUCTION

Cytoskeletal polymers are assembled and organized into specific structures to control cell shape, motility, and adhesion. While a great deal of research has identified a large number of proteins which regulate the assembly, dynamics, and organization of both microtubules and F-actin, very few proteins that regulate intermediate filament (IF) organization are known. Yet intermediate filaments are dynamic structures that rapidly respond to changes in a cell’s status (Windoffer et al., 2011). For example, IFs remodel during cell division, quickly form attachments with desmosomes and hemidesmosomes during the formation of these cell adhesion structures, assemble at the leading edge of migrating cells, and in response to externally applied forces on cadherins (Coulombe and Wong, 2004; Liao and Omary, 1996; Ridge et al., 2005; Weber et al., 2012; Windoffer et al., 2006). Further emphasizing the need to understand how IFs organize is the spectrum of diseases caused by mutations in IF genes, including blistering disorders in the epidermis caused by mutations in keratins and myopathies/cardiomyopathies associated with desmin mutations (Clemen et al., 2013; Eriksson et al., 2009; Smith, 2003).

The association of desmosome cell-cell adhesion complexes with IFs provides robust mechanical strength to tissues that experience force (Green and Simpson, 2007). Desmosome disorders of the skin encompass mild cases with skin thickening to lethal blistering (Kottke et al., 2006). Mutations in genes for desmosomal proteins also cause arrhythmogenic right ventricular dysplasia/cardiomyopathy – a heart defect characterized by fibro-fatty infiltrates (Delmar and McKenna, 2010). IFs associate with desmosomes through desmoplakin, a core component of the desmosomal plaque that directly binds to the epithelial keratin filaments and the desmin filaments in cardiac muscle (Bornslaeger et al., 1996; Gallicano et al., 1998; Kouklis et al., 1994; Stappenbeck et al., 1993). Until recently, there was no data that desmosomes actively control IF assembly or organization beyond simply binding pre-existing filaments. However, Leube and colleagues recently imaged events that appear to be the *de novo* nucleation of keratin filaments at desmosomes (Moch et al., 2020). This finding extended prior work demonstrating that keratin filaments were nucleated near focal complexes at the leading edge of migrating cells (Windoffer et al., 2006). In neither case is the molecular machinery required for this assembly known. These findings are intriguing as other cytoskeletal organizing structures – including centrosomes and adherens junctions – also influence the assembly and dynamics of the polymers they bind (Chhabra and Higgs, 2007; Kollman et al., 2011). Centrosomes recruit γ-tubulin to nucleate microtubules, while both the Arp2/3 complex and formins have been reported to promote local actin assembly at adherens junctions, yet no assembly factor has been identified for IFs (Kobielak et al., 2004; Moritz et al., 1995; Tang and Brieher, 2012; Verma et al., 2012; Zheng et al., 1995).

At present there is only a limited repertoire of proteins known to control keratin organization. These include proteins that bind to pre-formed filaments, like desmoplakin, as well as other plakin family members, e.g. plectin, which can crosslink keratins with other cytoskeletal structures (Sonnenberg and Liem, 2007; Suozzi et al., 2012). In addition, keratin assembly status can be controlled by post-translational modifications by kinases, glycotransferases, and other enzymes (Omary et al., 1998). This lack of understanding of polymer assembly is largely true for other IF proteins as well. However, the assembly of neurofilaments, an IF polymer that is quite distinct from keratins, is controlled by Ndel1 – a protein whose function has been best studied in neurodevelopment and in its association with Lis1 and dynein (Chansard et al., 2011; Nguyen et al., 2004; Niethammer et al., 2000). In neurons, Ndel1 was required for neurofilament organization. Strikingly, it directly promoted the assembly of neurofilaments from purified subunits (Nguyen et al., 2004). In addition, Ndel1 has been implicated in controlling the organization of both nuclear lamins and vimentins, two other IF proteins, although these effects have been attributed, at least in part, to dynein-directed transport (Ma et al., 2009; Shim et al., 2008).

In the epidermis, Ndel1, and its binding partner Lis1, localize to centrosomes in epidermal progenitor cells (Sumigray et al., 2011). Upon differentiation, both of these proteins relocalize to desmosomes where Lis1 is necessary to control desmosome-dependent microtubule reorganization (Lechler and Fuchs, 2007; Sumigray et al., 2011). In addition, loss of Lis1 resulted in desmosome defects, though Lis1’s role in desmosome stability is unknown. What role Ndel1 provides at the desmosome, and whether it is important for either microtubule or keratin filament organization, has not been addressed. Here we demonstrate that Ndel1 is required for proper desmosome function both in intact epidermis and in cultured keratinocytes. Ndel1 directly interacts with keratin subunits and promotes their local assembly/organization within the cell. Thus, we propose that desmosomes actively participate in keratin assembly and organization through the recruitment of Ndel1 to the cell cortex.

## RESULTS

### Ndel1 directly interacts with keratin subunits but not filaments

To begin to test the possibility that Ndel1 binds to keratins, we purified GST-tagged Ndel1 as well as keratin 10 (K10) from bacterial lysates. Ndel1 contains an N-terminal coiled-coil which is required for its interaction with Lis1 and contains an alpha-helix in its C-terminus that is important for interacting with a number of proteins, including DISC1 and dynein/dynactin (Fig 1A) (Soares et al., 2012). K10, together with keratin 1 (K1), are the most abundant keratins in the differentiated cells of the epidermis in which Ndel1 is localized to the desmosomes. With purified proteins we saw interaction of the full-length Ndel1 protein, as well as the isolated alpha-helix (AH) with K10 (Fig 1B). Using this binding assay, we performed structure/function analysis of K10 to define region(s) of the protein required for interaction. These studies demonstrated that the IF consensus motif of K10 was both necessary and sufficient for interaction with Ndel1 (Fig 1C). This short sequence is found in all intermediate filament proteins, suggesting that Ndel1 is likely to have direct interactions within the broad IF protein family (Fig 1, Supplement 1)(Geisler et al., 1983; Geisler and Weber, 1982). We performed mutagenesis across this conserved IF motif to identify important amino acids for Ndel1 binding. A conservative change of leucine 442 to alanine was sufficient to disrupt the interaction between Ndel1 and keratin 10 (Fig 1D). This leucine shows absolute conservation in the consensus motif across both vertebrate and invertebrate IFs (Fig 1, Supplement 1). Nde1, a paralog of Ndel1, showed similar ability to bind to K10 (Fig 1, Supplement 2). In addition, consistent with binding to the IF consensus sequence, it was able to interact with both type I and type II keratins including K1 and K5 (Fig 1, Supplement 2).

**Figure 1:**
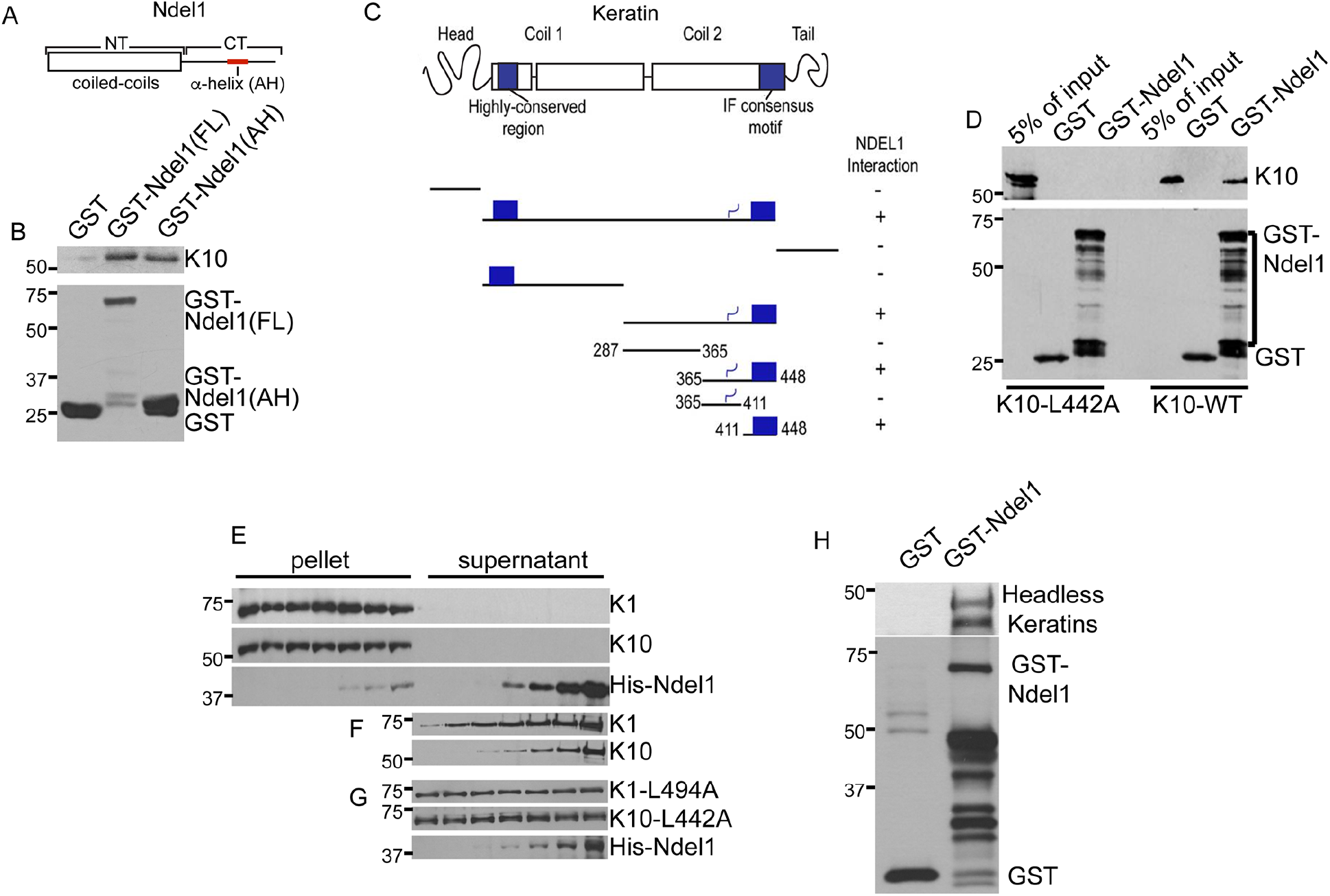
Ndel1 directly binds to keratin subunits. (A) Diagram of the domains of Ndel1. (B) GST-pull down assay showing interactions between Ndel1 and Keratin 10 (K10). Purified proteins were used to demonstrate a direct interaction between GST-Ndel1, both full length (FL) and the alpha-helical region (AH), and keratin 10. (C) Diagram of keratin domain structure and truncation mutants tested for keratin 10 interaction. Plus marks indicate Ndel1 interaction. Note that the IF consensus motif is sufficient for interaction. (D) Full-length keratin 10 or a point mutant, keratin 10(L442A) were mixed with glutathione-agarose beads bound to GST or GST-Ndel1. Specific interaction of WT but not mutant keratin was noted with GST-Ndel1. Note that the multiple bands in the GST-Ndel1 lane are degradation products. (E) Purified Keratin 1 and 10 were mixed in equimolar amounts and urea was step-wide dialyzed away to allow filament formation. After overnight incubation with increasing concentrations of His-Ndel1, the mixtures were ultracentrifuged. Pellet and supernatant were then analyzed by western blot. (F) A gel with the same samples loaded as in (E) but exposed for a longer time to visualize the levels of keratins in the supernatant. (G) The same experiment as in (E,F) except that mutant keratins that cannot interact with Ndel1 (K1-L494A and K10-L442A) were used. Note that there is no increase in soluble mutant keratin levels with increasing amounts of Ndel1. (H) Purified headless K1 and K10 (K1 amino acid 185-637 and K10 amino acids 129-561) were allowed to interact and then mixed with glutathione agarose bound to either GST or GST-Ndel1. A pan-keratin antibody detected association of both headless keratins with GST-Ndel1.

Keratin filaments form from obligate heterodimers of type I and type II keratins, such as K1/10 in the differentiated epidermis (Hatzfeld and Weber, 1990). Therefore, the interaction of Ndel1 with the keratin monomers described above is likely non-physiological, as most keratin exists in polymer form intracellularly, with significant pools of non-filamentous polymers – mostly tetramers – in addition to the fully polymerized IFs (Feng et al., 2013). We therefore tested the ability of Ndel1 to interact with keratin that had assembled into filaments. Equimolar amounts of keratin 1 and 10 in 8M urea were mixed and assembled into filaments by slowly dialyzing the urea away (Steinert and Gullino, 1976). Increasing amounts of Ndel1 were added to the pre-assembled filaments, allowed to bind overnight, and then pelleted by ultracentrifugation. Only very small amounts of Ndel1 were pelleted with the filaments, suggesting a low affinity between Ndel1 and filamentous keratin (Fig 1E). In contrast, when we overexposed gels with the soluble pool of keratins, we found that increasing Ndel1 levels resulted in increases in soluble, non-filamentous keratin subunits (Fig 1F). Upon mutation of each of the conserved leucines to alanines in the consensus motifs of keratin 1 and 10 (K1-L494A and K10-L442A), we did not observe this effect, demonstrating that Ndel1 must bind to keratin subunits to have this effect *in vitro* (Fig 1G). These data suggest that Ndel1 binds to soluble keratin subunits, but poorly to assembled filaments.

To gain additional information into the Ndel1/keratin interaction, we purified headless keratins which are unable to polymerize into filaments, but can still associate into soluble dimers and tetramers – two physiological species of IF subunits (Hatzfeld and Burba, 1994). Upon centrifugation, the majority of these proteins remained in solution (data not shown). When mixed with GST-Ndel1, these subunits were able to interact, demonstrating that Ndel1 is likely to interact with physiologically relevant keratin subunits (Fig 1H). Subunit binding proteins can have diverse effects on polymer assembly. They can sequester monomers, inhibiting polymerization, as is the case for thymosin β-4 for actin and stathmin for tubulin (Belmont and Mitchison, 1996; Goldschmidt-Clermont et al., 1992). Alternatively, they can act to promote polymerization (formins, profilin, and VCA domains in F-actin assembly, TOG domains in tubulin assembly (Firat-Karalar and Welch, 2011; Gunzelmann et al., 2018)). Therefore, we sought cellular assays to determine how Ndel1 might affect polymer dynamics and organization.

### Ndel1 affects keratin organization in keratinocytes

In order to more fully understand the effects that loss of Ndel1 has in cells, we established stable Ndel1 null keratinocytes (described in more detail below) along with wild-type (WT) controls from the back skin of postnatal day 0 (p0) mice. Immunofluorescent analysis of keratin revealed an increase in the perinuclear localization of keratins in Ndel1 null cells and a decrease in cortical filaments when compared to keratin networks in WT cells (Fig 2A,B). We further saw that WT cells included numerous filaments that extended well into the cell periphery which were less common in Ndel1 null cells. In addition, we examined the effect of expression of a GFP-tagged carboxy-terminal fragment of Ndel1 (Ndel1-CT-GFP) that lacks desmosome targeting activity but retains its keratin-interaction domain. Ndel1-CT-GFP did not localize to desmosomes and caused disruption of the keratin filament network (Fig 2C). Although it was able to change the organization of keratin filaments, we never saw Ndel1-CT-GFP (or full length Ndel1) colocalize with them. This observation is in line with our biochemical assays in Figure 1E which suggest Ndel1 has a low affinity for assembled IFs. In contrast, an amino-terminal fragment of Ndel1 that lacks its keratin-interaction domain (Ndel1-NT-GFP) localized to desmosomes but did not disrupt keratin organization appreciably (Fig 2D).

**Figure 2.**
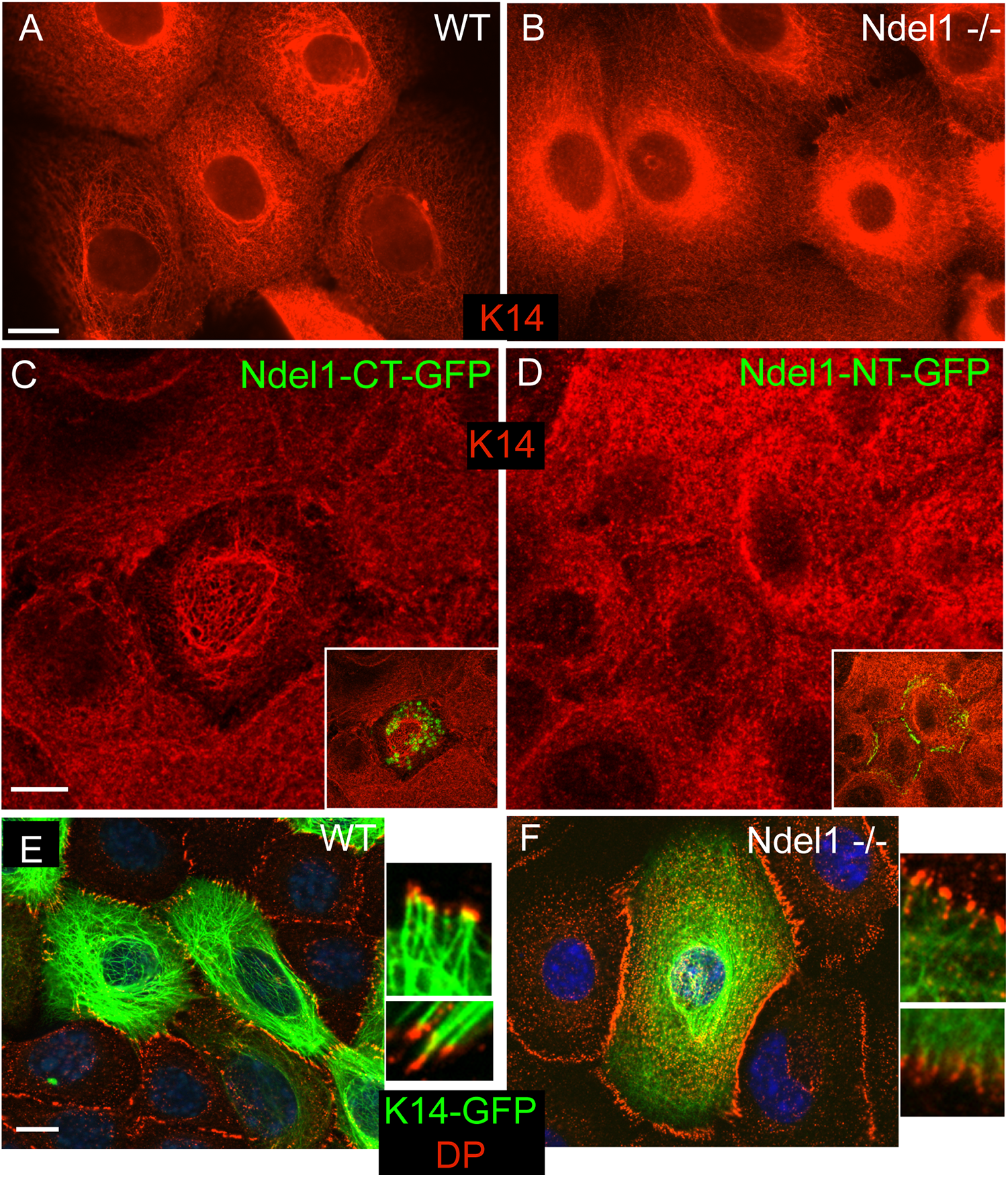
Ndel1 regulates keratin organization. (A,B) Endogenous keratin organization in WT and Ndel1 KO cells stained with anti-K5/14 antibodies. Cells were switched to high calcium-containing medium for 4 hours before fixation. (C) WT cells were transfected with Ndel1-CT-GFP and stained for keratin 14 (red). Note the partial collapse of the keratin network. (D) WT cells were transfected with Ndel1-NT-GFP and stained for keratin 14 (red). (E,F) WT and Ndel1 null cells were transfected with keratin 14-GFP (green), grown to confluency, switched to high calcium-containing media for 4 hours, and then fixed and stained for DP (red). Note the higher levels of peripheral keratin filaments and increased desmosome-keratin attachment in WT cells as compared to Ndel1 null cells. Scale bars are 10 μm.

We next transfected keratinocytes with either WT keratin 10 tagged with GFP (K10-GFP) or keratin 10 containing the mutation which we identified in Figure 1D that disrupts keratin 10’s interaction with Ndel1 (K10(L442A)-GFP). We found that there was an increase in the number of cells with abnormal keratin organization (16% with WT K10, 80% with K10-L442A - Fig 2, Supplement 1). In general, keratin aggregates rather than filamentous networks were observed in the mutant. This is consistent with Ndel1 interaction being important for proper keratin assembly.

We next asked whether Ndel1 promotes keratin organization at desmosomes, as Ndel1 localizes to these structures and they are important sites for keratin organization and assembly in keratinocytes (Moch et al., 2020; Sumigray et al., 2011). We therefore transfected cells with GFP-tagged keratin 14 (K14-GFP) to visualize keratin organization. After 4 hours in Ca^2+^-containing media, clear and strong connections between well-organized keratin filaments and desmosomes were observed at the desmosome in WT cells (Fig 2E). In Ndel1 null cells, while some keratin attachment was noted, there were fewer filaments at the cortex and the keratin network was poorly organized, as well as possessing the same increase in perinuclear localization we previously observed (Fig 2F). These data are consistent with Ndel1’s keratin binding activity playing an important role at the desmosome to promote keratin organization and attachment.

### Ndel1 promotes local keratin filament assembly

Our data suggest that Ndel1 could be acting at desmosomes to locally promote keratin filament assembly/organization, which we hypothesize is required for efficient attachment of desmosomes to the keratin network. This is supported not only by our biochemical and cell biological data, but also by the published finding that Ndel1 can promote neurofilament assembly kinetics (Nguyen et al., 2004). Unfortunately, the biochemical nature of keratins has prevented us from performing similar kinetic assays for assembly. To test whether Ndel1 is able to affect the local organization of keratins within the cell, we have taken a mistargeting approach. We fused Ndel1 to a mitochondrial membrane targeting sequence (mito-Ndel1) and coexpressed it with K14-GFP to report on keratin organization in wild-type keratinocytes. When a control construct of mito-PAGFP was transfected, keratin organization appeared normal and there was little overlap between mitochondria and the keratin network (Fig 3A). In contrast, expression of mito-Ndel1 induced a dramatic reorganization of the keratin network. Most prominently, we noted small structures in the cell that were enriched in K14-GFP and that were stained by the mitochondrial dye Mitotracker-Red (Fig 3B). Therefore, Ndel1 generated a local change in keratin organization when targeted to mitochondria. Upon extended incubation times after transfection, we noted defects in mitochondrial morphology (Fig 3C). Mitochondria became larger and either round or elongated tubules, and also showed a high concentration of keratin filaments around them, which is likely responsible for these morphological changes. However, it was difficult to discern true filamentous structures around the mitochondria in many cells due to the local concentration of keratins. To determine whether the keratins associated with these mitochondria were stable, similar to filamentous keratin, we performed fluorescence recovery after photobleaching (FRAP) on these cells. In both control and mito-Ndel1 cells, K14-GFP recovery was very slow, consistent with the majority of the mitochondrially-associated keratin being in its full-assembled, filamentous form (Fig 3D, E). This suggests that Ndel1 promotes the local assembly/organization of keratin filaments at specific subcellular sites.

**Figure 3.**
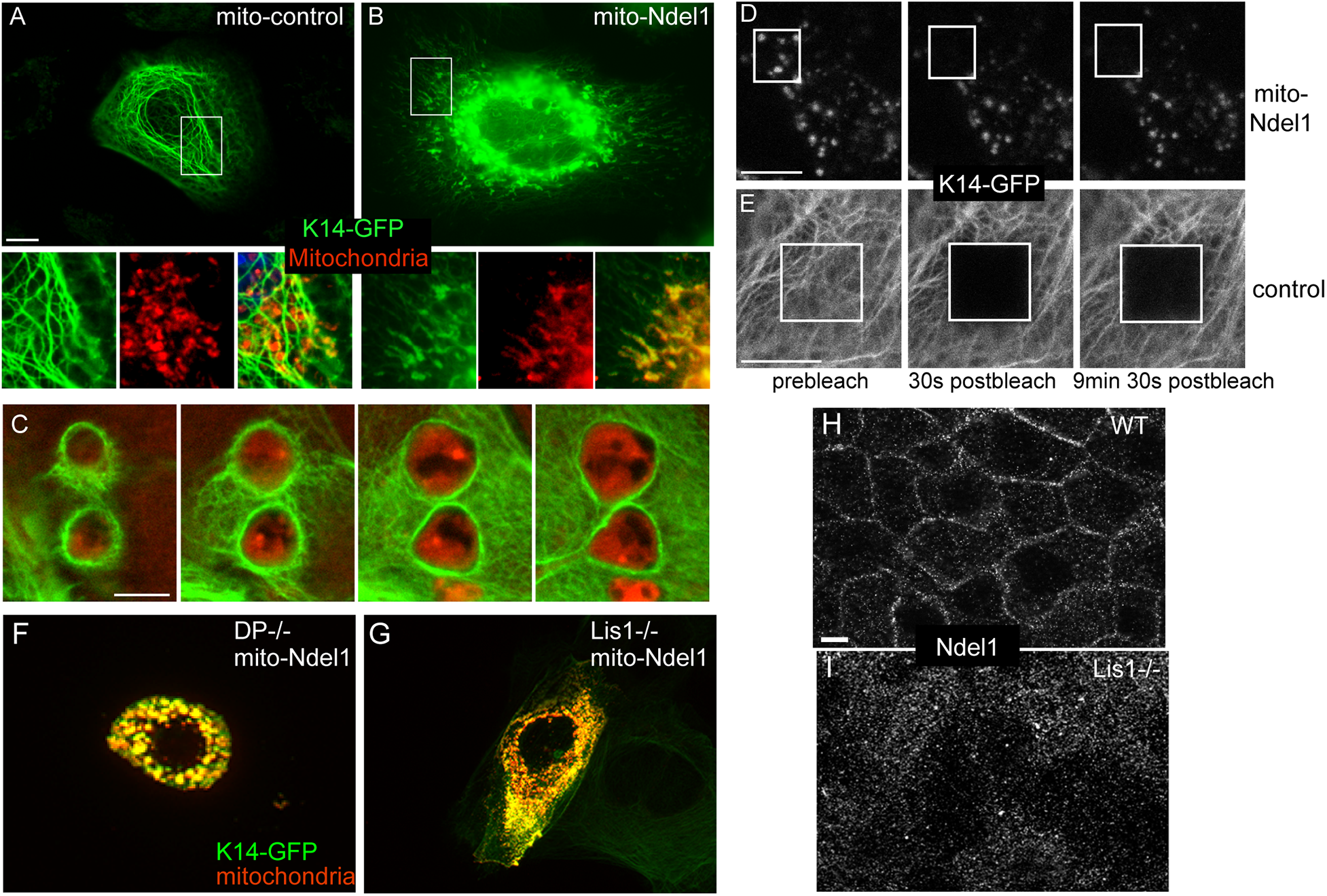
Ectopic Ndel1 induces keratin reorganization around mitochondria. (A,B) WT keratinocytes were co-transfected with keratin 14-GFP (green) and mito-control or mito-Ndel1 constructs. Mitochondria were labeled by incubating cells with MitoTracker-Red for 30 minutes before fixation. mito-Ndel1 induced a reorganization of keratins onto the surface of mitochondria. Scale bar is 10 μm. (C) Z-stack through enlarged mitochondria surrounded by keratin filaments. Red – mitochondria, Green – keratin 14-GFP. Scale bar is 5 μm. (D,E) FRAP analysis of keratin 14-GFP in control cell and cells transfected with mito-Ndel1. Scale bars in D,E are 10 μm. (F) DP null keratinocytes were transfected with mito-Ndel1 and keratin 14-GFP (green). Mitochondria were labeled with MitoTracker Red. (G) Lis1 null keratinocytes were transfected with mito-Ndel1 and keratin 14-GFP (green). Mitochondria were labeled with MitoTracker Red. (H,I) Immunofluorescence of Ndel1 in WT and Lis1 null cells. Scale bars are 10 μm.

### Desmoplakin and Lis1 are required for Ndel1 targeting, but not keratin filament assembly

One trivial explanation for Ndel1’s effects on keratin organization could be that it recruits desmoplakin to mitochondria. To test this, we examined the localization of GFP-tagged desmoplakin (DPI-GFP) in mito-Ndel1 expressing cells. We were never able to visualize any localization of DPI-GFP at mitochondria in these experiments (Fig 3, Supplement 1). To directly test whether desmoplakin is required for Ndel1’s keratin reorganization activity, we expressed mito-Ndel1 in desmoplakin null keratinocytes. These cells showed no defect in keratin accumulation around mitochondria, demonstrating that desmoplakin is not required for this activity (Fig 3F).

Lis1 binds to Ndel1 and its loss results in desmosome defects (Sumigray et al., 2011). We therefore wanted to test whether Ndel1’s keratin assembly activity required Lis1. We expressed full-length mito-Ndel1 in Lis1 null cells. To perform this experiment, Lis1 was ablated in Lis1 flox/flox cells by infection with high-titer adenoviral-Cre recombinase. We waited 24 hours after infection before transfection of the mito-Ndel1 construct to ensure complete loss of Lis1 (Sumigray et al., 2011). K14-GFP localized around mitochondria in these cells, demonstrating that Lis1 is not required for mito-Ndel1’s effects on keratin organization (Fig 3G). In addition, because Lis1 and Ndel1 interact with dynein and regulate its activity, we tested the roles of both microtubules and dynein in keratin assembly on mitochondria. Neither nocodazole treatment, which disrupts microtubules, nor treatment with the dynein inhibitor, ciliobrevin D, affected keratin assembly (Fig 3, Supplement 1). Therefore, Ndel1 is unlikely to act via localized transport of keratin subunits to specific cellular sites.

The above data demonstrated that neither desmoplakin nor Lis1 was required for Ndel1’s keratin assembly activity. It did, however, raise the question of why loss of Lis1 resulted in desmosome defects (Sumigray et al. 2011). We therefore asked whether Lis1 might function to target Ndel1 to the desmosome in the same way that desmoplakin is required for Ndel1 localization. Analysis of Ndel1 by immunofluorescence in WT and Lis1 null cells revealed an essential role for Lis1 in recruiting Ndel1 to the cell cortex (Fig 3H,I).

Both desmoplakin and Lis1 are required for the cortical reorganization of microtubules that occurs upon microtubule stabilization (Lechler and Fuchs, 2007; Sumigray et al., 2011). Ndel1 binds to Lis1 and has been demonstrated to control microtubule organization in other cellular contexts (Guo et al., 2006). We therefore tested whether Ndel1 was also necessary for the desmosome-dependent reorganization of microtubules into cortical arrays and whether this might underlie the desmosome defects. WT and Ndel1 null cells were treated with DMSO or with taxol (5 μM, 1h) to stabilize microtubules and induce their reorganization. We observed a robust reorganization of microtubules to the cell cortex in both WT and Ndel1 null cells upon taxol treatment (Fig 3, Supplement 2). These data demonstrate that the keratin and microtubule regulating functions of this complex are separable.

### Ndel1 is essential for desmosome stability

Having demonstrated that Ndel1 is capable of affecting local keratin organization, we next looked at the consequences of Ndel1 depletion on desmosomes. Desmosomes are not only the major organizers of the keratin filament network in the differentiated epidermis, but they also depend on bound keratins for their stability and function. We examined the localization of desmosomal proteins in both control and Ndel1 null keratinocytes. The localizations of the core desmosome components, plakoglobin and plakophilin 2, were decreased at cell-cell junctions and their cytoplasmic pools were increased (Fig 4A-D). Though perturbation of desmoplakin localization was less dramatic, it too showed a decrease in cortical staining and an increase in the cytoplasmic pool (Fig 4E, F). When we examined the total levels of desmosomal proteins in epidermal lysates, we noted minor decreases in the amounts of several desmosomal proteins, including desmoplakin and plakophilin 2 (Fig 4G). Desmosomes are typically very insoluble in keratinocytes due in part to their interactions with the keratin filament network. When we isolated soluble and insoluble pools of desmosomal proteins, we observed a dramatic increase in the soluble pools of all of the desmosomal components tested (Fig 4H). Therefore, Ndel1 cells have perturbed desmosomes in cultured cells, as revealed by both cell biological and biochemical measures.

**Figure 4.**
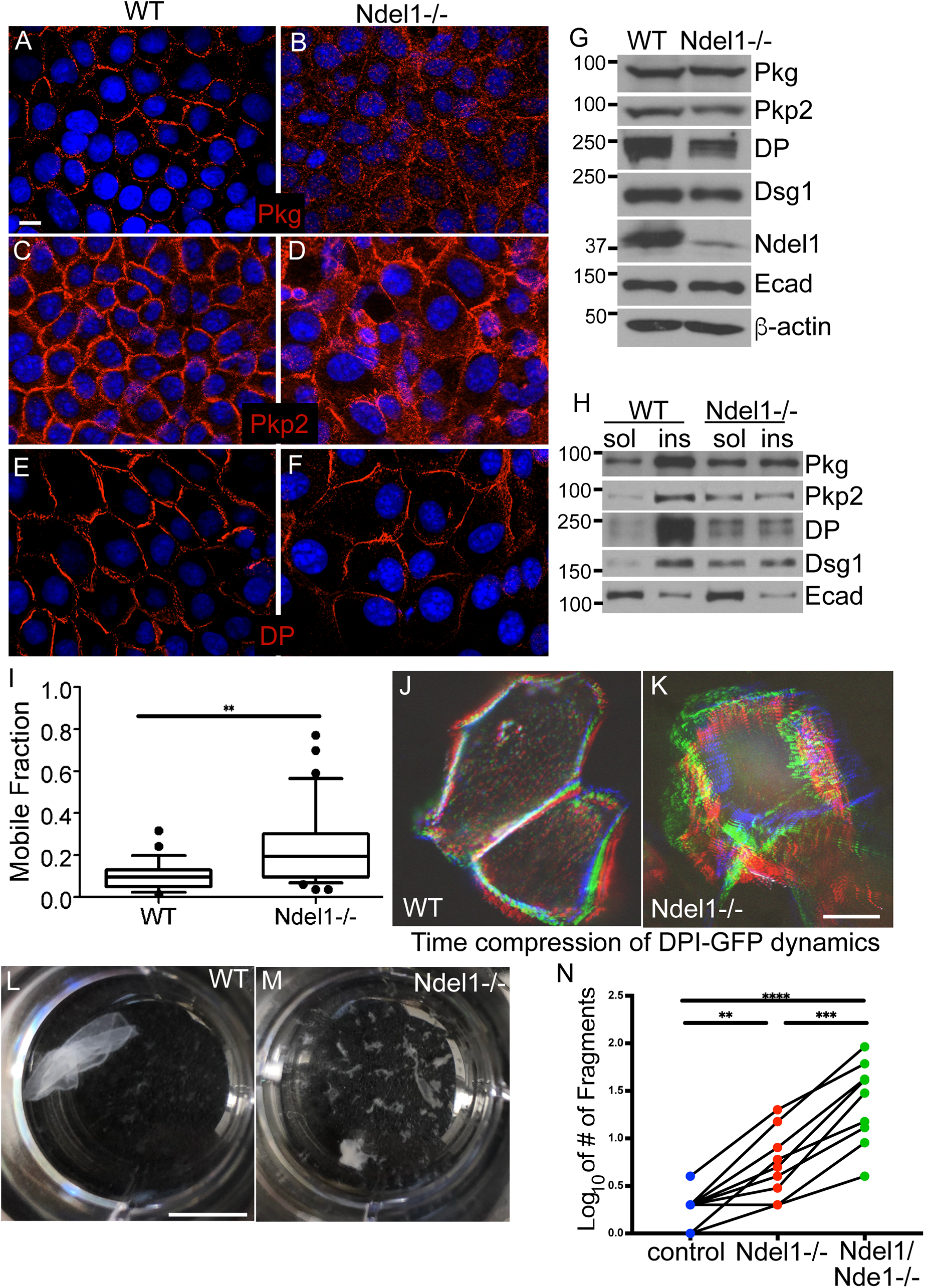
Ndel1 is required for desmosome stability. (A-F) Immunofluorescence analysis of desmosomal proteins in WT and Ndel1 null keratinocytes, as indicated. Scale bar is 10 μm. (G) Western blot analysis of desmosomal and adherens junction protein levels in lysates prepared from WT and Ndel1 null keratinocytes. (H) Protein lysates from control and Ndel1 null cells were fractionated into soluble and insoluble components. Proteins levels were then analyzed by western blotting. (I) Fluorescence recovery after photobleaching (FRAP) was measured for DPI-GFP transfected into WT and Ndel1 null cells. FRAP was performed on cells grown in high calcium-containing media for 24 h to ensure desmosome assembly. (J,K) Time-compressed stacks from Videos 1 and 2. Movies were aligned for drift, maximum pixel intensities are shown and the timepoints were color coded as RGB images. Note the increased dynamics of DPI-GFP in the Ndel1 null cells. Scale bar is 10 μm. (L,M) Dispase-based dissociation assays were done with confluent monolayers of keratinocytes incubated for 72 h after Ca^2+^ switch and treated with dispase. The lifted monolayers of keratinocytes were subjected to mechanical stress. Scale bar is 1 cm. (N) Quantitation of dispase assay. Data shown are from 8 paired experiments.

We next performed FRAP analysis of DPI-GFP in control and Ndel1 null keratinocytes to determine whether desmosome dynamics as well as solubility were affected by loss of Ndel1. Consistent with studies in Lis1 null keratinocytes (Sumigray et al., 2011), we found that there was an increased mobile fraction, suggesting that desmosomes are less stable in Ndel1 null keratinocytes (Fig 4I). In addition, we performed time-lapse imaging of these cells 24 hours after calcium induction of desmosome formation. WT cells consistently showed very stable membranes, with little movement (Supplementary Video 1). In contrast, Ndel1 null cells showed highly dynamic cell-cell adhesions that changed considerably over the 2 hours of imaging (Supplementary Video 2). Time-lapse compressions were generated and color-coded to demonstrate the changes in cell shape and desmosome positioning (Fig 4J, K).

The above data demonstrated that the stability of desmosomes was disrupted in the Ndel1 null keratinocytes. The adhesive activity of keratinocytes is largely due to the presence of robust desmosomes (Vasioukhin et al., 2001). To functionally test the strength of desmosomal adhesions, confluent sheets of keratinocytes were detached from the plate on which they were grown by treatment with dispase. The sheets were then mechanically disrupted and the number of cell fragments that resulted were counted. While WT cells were quite mechanically robust, Ndel1 null cells were notably fragile and fragmented into more pieces, consistent with a defect in desmosomal adhesions (Fig 4L-N).

Ndel1 has a paralog, Nde1, that has high homology and also localizes to desmosomes (Soares et al., 2012; Sumigray et al., 2011). To test whether Nde1 might have partially redundant functions with Ndel1, we used CRISPR/Cas9 genome editing to knock out Nde1 in Ndel1 null cells. Western blotting confirmed the loss of protein (Figure 4, Supplement 1). These mutant cells showed an enhanced mechanical fragility as compared to the control as well as the Ndel1 null cells (Fig 4N). Staining for desmosomal proteins revealed similar changes in localization as described above for Ndel1 null cells (Figure 4, Supplement 1). Comparable results were found for a second clone with independent mutations (data not shown). Together these data demonstrate that Ndel1, and its paralog Nde1, are desmosome-associated proteins that are required for desmosomes’ adhesive function.

### Ndel1 depletion in the epidermis leads to skin defects

To test the functional role of Ndel1 at desmosomes *in vivo*, we have generated mice in which Ndel1 is conditionally lost in the epidermis through recombination using Cre recombinase under the control of the keratin 14 promoter (referred to as Ndel1 cKO) (Vasioukhin et al., 1999). Cre expression begins around e14.5 in epidermal progenitors, and at birth, Ndel1 cKO pups had dramatically reduced levels of Ndel1 in epidermal lysates (Fig 5A). Unlike ablation of its binding partner, Lis1, loss of Ndel1 in the epidermis did not result in neonatal lethality (Sumigray et al., 2011). This may be due, in part, to the presence of Nde1 which has overlapping functions with Ndel1 in cultured keratinocytes (Fig 4N).

**Figure 5.**
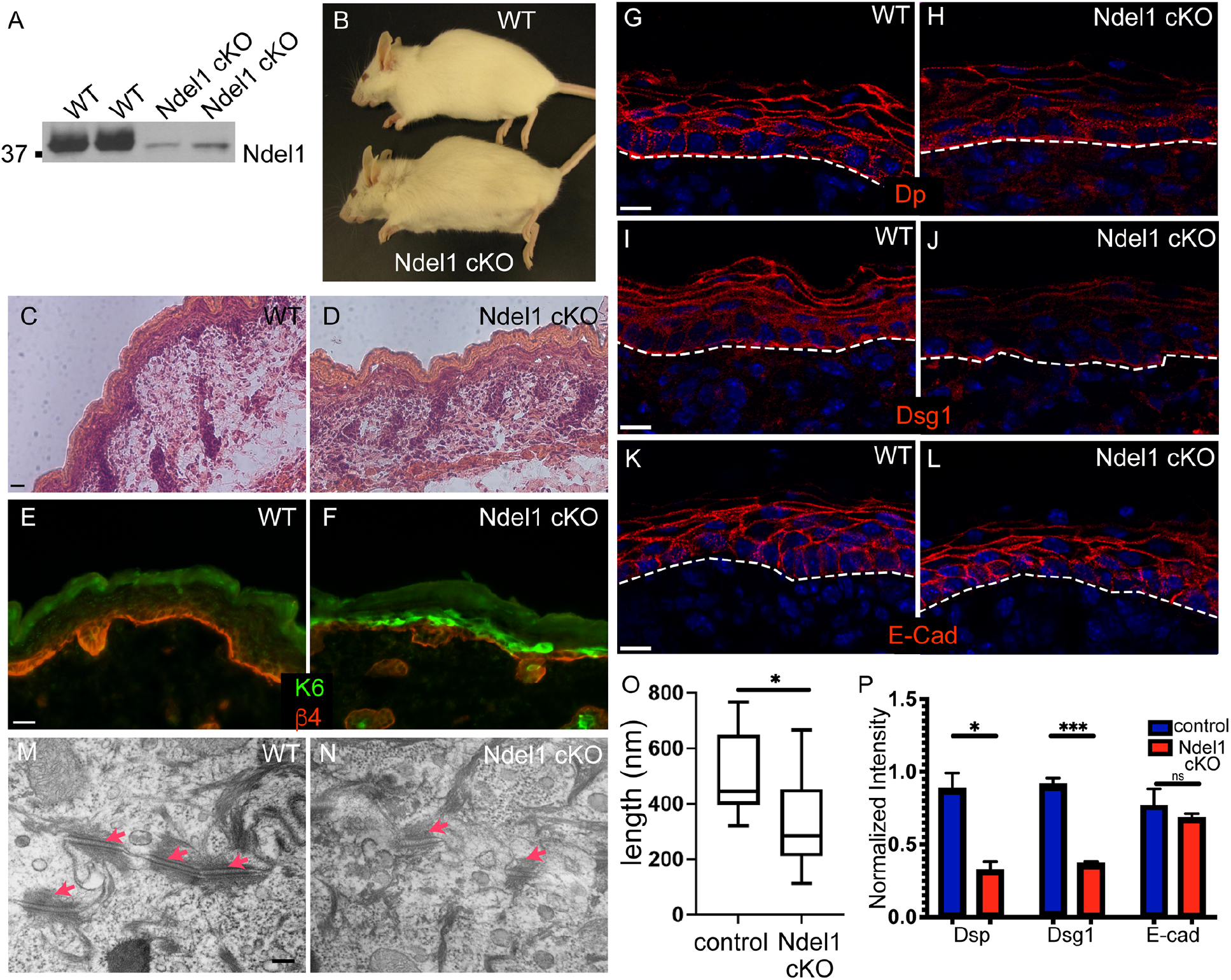
Loss of Ndel1 in the epidermis results in desmosome defects. (A) Western blots of epidermal lysates from 2 wildtype (WT) and 2 Ndel1 cKO neonatal mice. (B) Photo of five month old WT and Ndel1 cKO mice. (C,D) Hematoxylin and eosin stained sections from WT and Ndel1 cKO p0 mice. (E,F) Staining for the stress marker keratin 6 (green) and the basement membrane marker β-4 integrin (red) in WT and Ndel1 cKO p0 skin. (G-L) Staining of desmosomal and adherens junction components in WT and Ndel1 cKO p0 skin sections as indicated. All scale bars are 10 μm. (M,N) Transmission electron microscopy of spinous cells in control and Ndel1 null epidermis. Red arrows indicate desmosomes. (O) Quantitation of desmosome length in control and Ndel1 null epidermis. (P) Quantitation of cortical levels of desmoplakin (Dsp), Dsg1, and E-cadherin in control and Ndel1 null epidermis.

We were unable to reliably identify Ndel1 cKO pups at birth by any gross phenotype. However, adult Ndel1 cKO mice were identifiable by a ragged appearance to their hair coat (Fig 5B). Histologic analysis of p0 pups demonstrated a relatively normal looking stratified epidermis (Fig 5C, D). There were some focal regions with micro-blisters and small cell-cell separations, though these were not prominent throughout the epidermis (Fig 5D). Ablation of some desmosomal genes, such as plakoglobin, also result in milder phenotypes than the blistering seen upon complete loss of desmoplakin (Li et al., 2012). Consistent with a mild effect, several observations demonstrated that the skin was adversely affected by the loss of Ndel1. This included an upregulation of the stress-induced protein keratin 6 in focal regions of the epidermis (Fig 5E, F). Because of Ndel1’s localization to desmosomes and our findings in cultured keratinocytes, we examined the status of these cell adhesion structures. Staining for desmoplakin and Dsg1 showed decreased junctional staining (Fig 5G-J, P). This effect was specific to desmosomes as normal localization of the adherens junction component E-cadherin was observed (Fig 5K, L, P). Because of the decrease in junctional desmosome components that we saw through immunofluorescence, we next took advantage of the electron dense nature of desmosomes and analyzed them via transmission electron microscopy (TEM). Strikingly, we found that the desmosomes in the epidermis of Ndel1 cKO mice were smaller and less electron dense than those in the control epidermis (Fig 5M,N). Quantification of mutant desmosomes revealed that they were roughly about half the length of the controls (Fig 5O). Our data thus support a model that Ndel1/Nde1 promote the local assembly/reorganization of keratin filaments at desmosomes, and, when these proteins are depleted, the lack of strong, stable attachments to keratin causes desmosome instability and leads to epidermal defects.

## DISCUSSION

Cells have developed many ways to control the local assembly and organization of cytoskeleton filaments. The identification of factors that promote assembly of microtubules and F-actin has allowed a much deeper understanding of how these cytoskeletal networks form and of the signaling pathways that govern local assembly. This regulation occurs through both localization and activation of these organizers at specific cellular sites. Our data suggest that Ndel1 promotes the local assembly of keratin filaments. Evidence in this paper as well as previously published data suggest that Ndel1 may be a general factor that promotes local IF assembly and organization across diverse cell types. In addition to Ndel1’s reported localization to desmosomes, where it is responsible for robust desmosome/keratin attachment (Sumigray et al., 2011), Ndel1 has also been reported to localize to nascent adhesions at the leading edge of migrating cells, sites where keratin filament nucleation has also been observed (Shan et al., 2009; Windoffer et al., 2006). More broadly, Ndel1 localizes to the mitotic spindle where it has been further implicated in nuclear lamin assembly/organization (Ma et al., 2009). Additionally, Ndel1 localizes to centrosomes, which have been a long appreciated foci for IF organization in some cell types (Kalnins et al., 1985; Niethammer et al., 2000; Sasaki et al., 2000). Thus, in diverse cell types and subcellular locales, Ndel1 promotes IF assembly/organization. Understanding the molecular mechanism of Ndel1, and its regulation by signaling pathways, are clearly important future areas of investigation.

Two models have been proposed to explain Ndel1’s role in IF assembly. First, Ndel1 could directly control the nucleation and/or elongation of IF filaments, as suggested by the finding that purified Ndel1 promotes neurofilament assembly *in vitro* (Nguyen et al., 2004). A second, and not mutually exclusive possibility, is that Ndel1 works in a more canonical Lis1/dynein-dependent manner to transport IF subunits or small filaments to distinct cellular locations. This has been proposed for both nuclear lamins and vimentin (Ma et al., 2009; Shim et al., 2008). While our data support the idea that Ndel1 can promote local assembly of keratins, there is currently no kinetic assay for keratin assembly *in vitro*, precluding a direct testing of the first model. However, we find that Ndel1 lacking both the Lis1 binding site and the amino-terminal dynein binding site is still functional in promoting keratin assembly when targeted to mitochondria. This region also disrupts normal keratin organization when it can’t localize to desmosomes. We did not find any effect on keratin assembly activity in cells devoid of Lis1 nor in cells treated with either nocodazole or dynein inhibitors. Therefore, our data are consistent with a more direct role for Ndel1 in keratin filament nucleation and/or elongation, similar to what has been reported for neurofilaments (Nguyen et al., 2004). Both structural data and structure/function analysis of Ndel1 and keratins should begin to define how Ndel1 affects keratin assembly. The fact that Ndel1 binds to the consensus motif which defines a conserved sequence present in all IF proteins is especially interesting. This region may thus have both important structural and regulatory roles in keratin assembly. Understanding the specific types of soluble keratin subunits that Ndel1 interacts with will also be important in determining how it promotes keratin assembly.

Ndel1 ablation resulted in desmosome defects in addition to keratin organization defects. While it is possible that Ndel1 has additional structural roles in desmosome stability, the desmosome defects are likely secondary to the keratin mis-organization. A number of studies have reported roles for keratins in desmosome formation and/or stability. For example, defects in desmosome structure have been reported in samples from human patients with desmoplakin mutations (Wan et al., 2004). In addition, keratin 8 controls the desmosomal localization of desmoplakin in hepatocytes, the ablation of keratin 1 and 10 in the epidermis resulted in smaller desmosomes, and the loss of all keratins led to desmosome defects in embryonic epithelia (Loranger et al., 2006; Vijayaraj et al., 2009). Our data further support an important role for desmosome-keratin attachments in regulating desmosome stability.

The loss of Ndel1 has a much less severe phenotype than loss of Lis1 or desmoplakin in the epidermis (Sumigray et al., 2011; Vasioukhin et al., 2001). Its severity is much more similar to loss of plakoglobin, which is not lethal when conditionally ablated in the epidermis (Li et al., 2012). This suggests a partial loss of desmosome function. In fact, our data demonstrate that the desmosomes are present and functionally able to direct microtubule reorganization, though they are deficient in robust keratin attachment. We demonstrated that Nde1, a homolog of Ndel1, acts in a somewhat functionally redundant manner. Nde1 shows a similar localization pattern as Ndel1 in keratinocytes and also binds to keratin subunits (Sumigray et al., 2011) data not shown). Future genetics approaches are needed to determine the phenotype of combined Ndel1/Nde1 loss on the epidermis.

While the desmosome has often been portrayed as a rather passive structure, our data highlight its active role in controlling both keratin and microtubule organization. Strikingly, these are both mediated, in part, by the recruitment of a complex of proteins containing Lis1 and Ndel1 to the desmosomes.

## MATERIALS AND METHODS

### Mice

K14-Cre (Vasioukhin et al., 1999) and Ndel1 fl/fl (Sasaki et al., 2005) have been described. All animal work was approved by Duke University’s Institutional Animal Care and Use Committee.

### Keratinocyte cultures

Ndel1 null keratinocytes were isolated from back skins of newborn pups. All keratinocytes were maintained in low Ca^2+^ (0.05 mM Ca^2+^) E media containing 15% fetal bovine serum (Hyclone) in a humidified incubator (37°C and 7.5% CO_2_). For microtubule organization assays, cells were treated with taxol after induction of differentiation for 1 h at 10 μM. Nocodazole (10 μM, Sigma) and ciliobrevin D (50 μM, Millipore) were added to cells for 1 hour before fixation. MitoTracker Red (Invitrogen) was added to cells as per manufacturer’s instructions. All transfections were performed with Mirus TransIT transfection media according to manufacturer’s protocols. Cell adhesion was induced by addition of calcium (1.2mm) to the media.

### Immunofluorescence staining

Keratinocytes were grown to confluence on glass coverslips in low Ca^2+^ media until confluent. After 24 h under terminal differentiating conditions (1.2 mM CaCl_2_, where appropriate), cells were fixed with either 4% PFA at 37°C for 10 minutes or in methanol for 2 min at −20°C. After fixation, cells were washed with PBST and then blocking buffer containing 5% BSA, 5% normal donkey serum and 5% normal goat serum. Staining of cryosections of back skin was performed in the same way as explained above. Imaging was performed on an AxioImager Z1 microscope with Apotome attachment and MRm camera (Carl Zeiss). A 63X, 1.4NA Plan Apotchromat objective and Immersol 518F immersion oil were used (Carl Zeiss). Imaging was performed at room temperature through either 12 mm round coverslips (No. 1) for cells, or No. 1.5 coverslips (VWR) for tissue sections. The following primary antibodies were used: rabbit anti-Ndel1 (Sumigray et al., 2011), mouse anti–β-tubulin and β-actin (Sigma-Aldrich), rabbit anti-keratin 6 and rabbit anti-keratin 10 (Covance), anti-keratin 5/14 (generated in lab), mouse anti-DP1/2 (Chemicon), guinea pig anti-plakophillin2 (Progen), mouse anti-desmoglein1 and rat anti-CD104 (BD Biosciences), mouse anti-CD3 (Millipore), guinea pig anti-Dsc1 (gift from I. King, Medical Research Council, London, England, UK), rat anti-E-cadherin and rabbit anti-keratin 1 (gifts of C. Jamora, InStem, India).

### FRAP analysis

Sample preparation and FRAP setup were done as described previously (Foote et al., 2013). In brief, mouse keratinocytes were grown on 35-mm glass-bottom culture dishes (no 1.5; MatTek Corporation) and transfected with DPI-GFP (provided by K. Green, Northwestern University, Chicago, IL) using Mirus *Trans*IT®-keratinocyte transfection reagent. Calcium was added to cells to 1.2 mM and junctions were allowed to form for 24 hours before imaging. A Zeiss 710 Confocal scanning light microscope with 63X 1.4 NA objective and Zen software were used to determine the mobile fraction of desmoplakin at cell cortical regions. Imaging was performed at 37°C . Regions of interest were bleached and only those cells in which 70% or more bleach was accomplished were analyzed. The fluorescence intensities were calculated by MetaMorph image analysis software and normalized to background intensity. The mobile fraction was determined as mf = I_max_-I_o_/1-I_o_ (Shen et al., 2008). Three different experiments were performed and statistical analysis was performed with combined data.

### Dispase-based dissociation assay

Mouse keratinocytes in 12 well plates were grown in low Ca^2+^ media until confluent. After 48 h under differentiating conditions (1.2 mM CaCl_2_), cells were washed twice with PBS and incubated with 2.4 U/ml dispase (Roche) for 1h at 37°C. Floating monolayer sheets were pipetted using a cut p1000 pipette tip. Pictures of the resulting sheet fragments were taken with a Canon Powershot camera and the number of fragments was determined by ImageJ software using the cell counter function. Tukey’s post-hoc test was performed after RM-ANOVA detected significance with Graph Pad Prism8 software to determine significance.

### CRISPR-Cas9 Genome Editing

The gRNA against Nde1 was designed using E-CRISP (Heigwer et al., 2014), and inserted into the pSpCas9(BB)-2A-Puro vector as described in (Ran et al., 2013), resulting in the pSpCas9-puro-Nde1 construct. Keratinocytes raised in the conditions described above were transfected with the pSpCas9-puro-Nde1 construct using Mirus *Trans*IT® transfection reagent. 24 hours after transfection, cells were subjected to selection by treating with 2μg/mL puromycin for 72 hours. The surviving cells were allowed to recover without puromycin for 7 days before being clonally isolated via serial dilution. Once lines were established from the resultant clones, Nde1 levels were analyzed by western blotting and the targeted locus was sequenced to characterize the mutations.

### Statistics

Student’s T-test was used for pair wise comparisons. For the dispase assays, cell sheets from the same plate were treated as repeated measures and RM-ANOVA was performed on the data using Tukey’s post-hoc test for multiple comparisons to analyze the individual genotypes for significance. (*) signifies a p-value less than 0.05, (**) is one less than 0.01, (***) is less than 0.001, and (****) is a p-value less than 0.0001.

### Protein-Protein Interactions

GST fusion proteins were produced in bacterial cells upon induction with 1 mM IPTG. The *E. coli* were harvested and lysed by sonication for 3X 10 seconds in ice-cold lysis buffer containing 50 mM Tris-HCl, pH 7.4, 2 mM MgCl_2_, 50 mM NaCl_2_, 10% glycerol, 1% Triton-X, 1 mM DTT, 1 mM PMSF, and protease inhibitors. After centrifugation at 12,000 rpm for 10 min at 4°C, the supernatants were obtained and incubated with glutathione-Sepharose beads for 6h at 4°C, and then washed extensively with lysis buffer. Keratins were isolated from bacterial inclusion bodies, solubilized in 8M urea, and then dialyzed into PBS. GST fusion proteins (0.5 mg/ml) bound to beads were mixed with keratins (0.3 mg/ml). After washing, proteins on beads were eluted through the addition of sample buffer and boiling. Eluted proteins were analyzed by western blotting.

### *In vitro* sedimentation assay

Equimolar amounts of purified keratin pairs in buffer containing 8M urea were slowly dialyzed into PBS with 1 mM DTT. After pipetting up and down to suspend the filaments, the complexes (12 μg/ml) were evenly distributed to tubes in keratin assembly buffer containing 50 mM Tris (7.4 pH), 100 mM NaCl, 0.1 mM EDTA, 0.1 mM EGTA, 1 mM DTT and protease inhibitors with His-Ndel1 (at 0, 1, 2, 6, 12, 25, and 50 μg/ml final concentration), and incubated at 4°C overnight. The mixtures were loaded into ultracentrifuge tubes (7 x 20 mm, Beckman Coulter) and fractionated at 100,000Xg using a Beckman TL-100 ultracentrifuge. Supernatant was removed and the pellets which were then resuspended in the same volume of sample buffer, and analyzed by immunoblotting.

### Preparation of protein lysates

Keratinocytes were washed twice with ice-cold PBS and scraped into lysis buffer containing 50 mM Tris-HCl, pH7.5, 150 mM NaCl, 50 mM NaF, 1 mM sodium pyrophosphate, 1% NP-40, 0.1 % sodium deoxycholate. After incubation for 30 min on ice, lysates were separated into soluble and insoluble fractions by centrifugation at 15,000Xg for 10 minutes.

### Transmission electron microscopy

Skin was harvested from neonatal mice and in 2% glutaraldehyde, 4% PFA, 1 mM CaCl_2_, and 0.05 M cacodylate (pH 7.4), for 1 h at room temperature and then overnight at 4°C. Samples were washed in 0.1 M sodium cacodylate buffer containing 7.5% sucrose. Samples were postfixed in 1% osmium tetroxide in 0.15 M sodium cacodylate buffer for 1 h and then washed in two changes of 0.11 M veronal acetate buffer for 15 min each. Samples were placed into en bloc stain (0.5% uranyl acetate in veronal acetate buffer) for 1 h, washed in veronal acetate buffer, and then dehydrated in a series of 70, 95, and 100% ethanol. Finally, the samples were prepared for embedding in Epon resin. After sectioning, samples were imaged with a CM12 transmission electron microscope (Phillips, now part of FEI, Hillsboro, OR) run at 80 kV with an XR60 camera (Advanced Microscopy Techniques, Woburn, MA). Image acquisition was done using 2Vu software (Advanced Microscopy Techniques).

## Acknowledgements

We thank Shinji Hirotsune for Ndel1 floxed mice, Pierre Coulombe for keratin constructs, Henry Foote for assistance with imaging, analysis and statistics, and Julie Underwood for excellent care of the mice. We are grateful to members of the Lechler lab for valuable comments throughout this work, and on the manuscript. This work was supported by NIH-NIAMS grants (R01AR055926 and R01AR067203 to TL), and by a Basil O’Connor award from the March of Dimes.

---

**Figure 1, Supplement 1.**
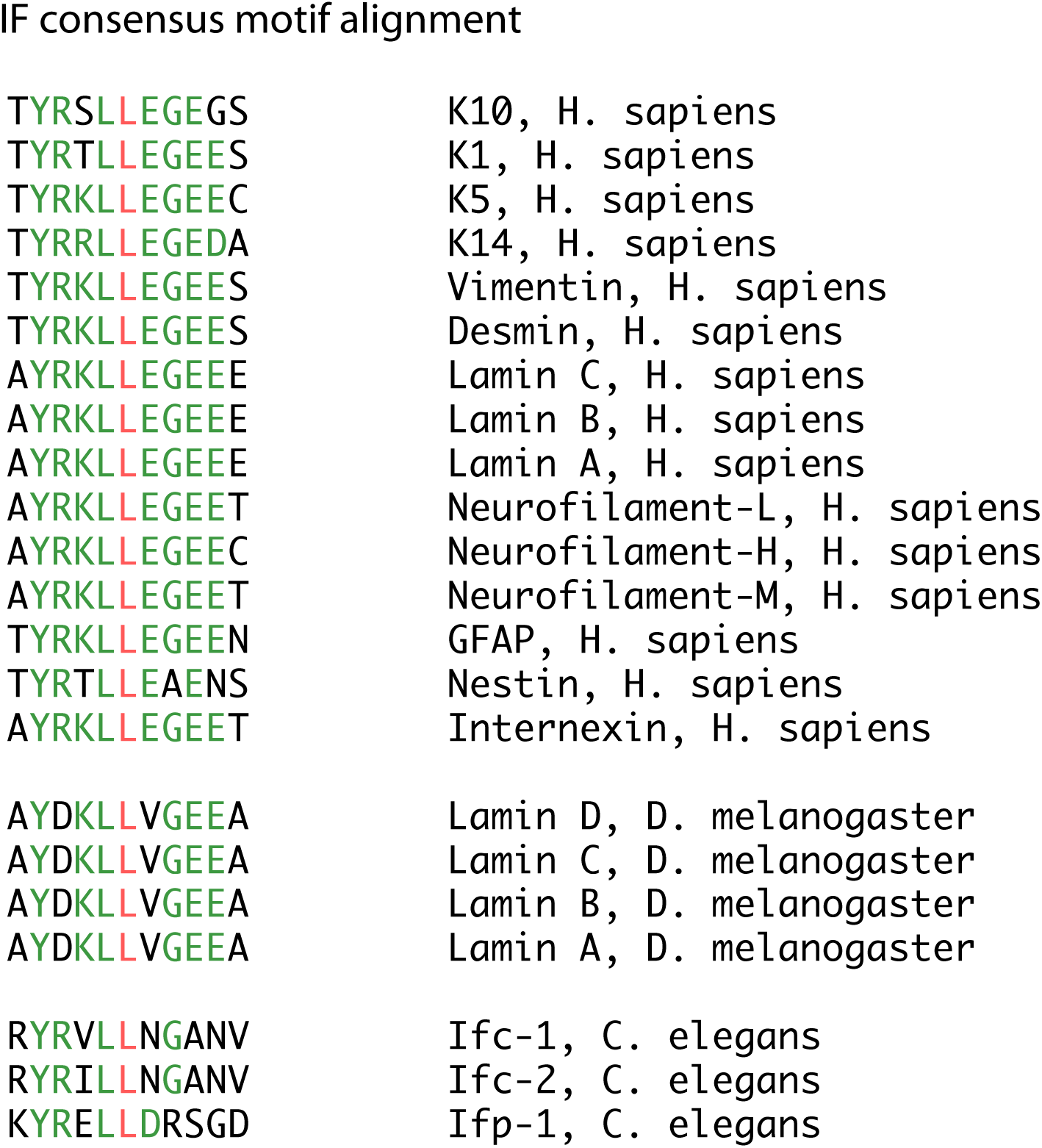
Alignment of intermediate filament (IF) consensus motifs of diverse IF proteins.

**Figure 1, Supplement 2.**
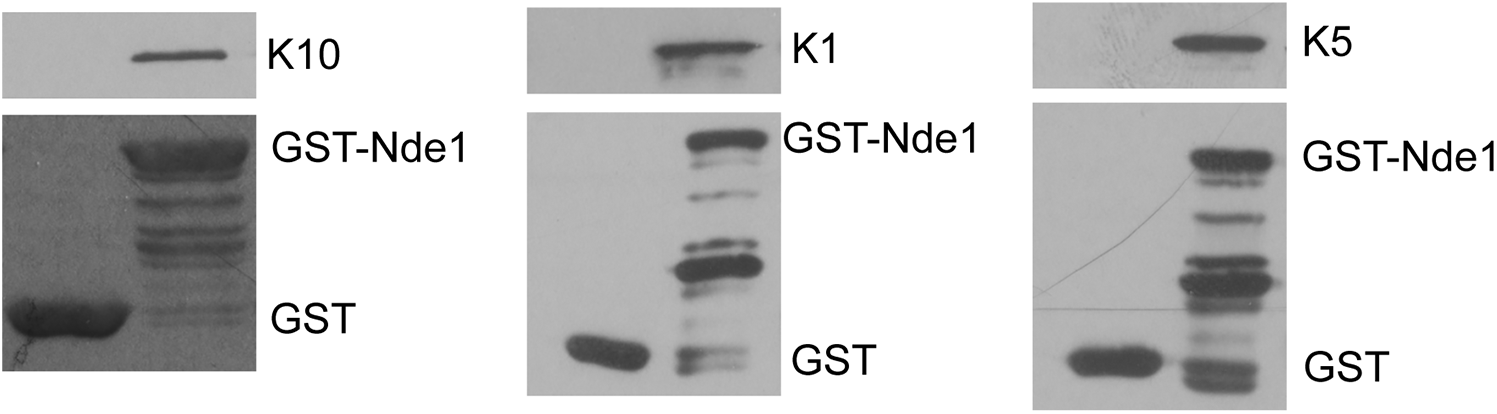
Interaction of Keratins 10, 1, and 5 with Nde1. Purified K10, K1, or K5 was added to beads bound to either GST or GST-Nde1.

**Figure 2, Supplement 1.**
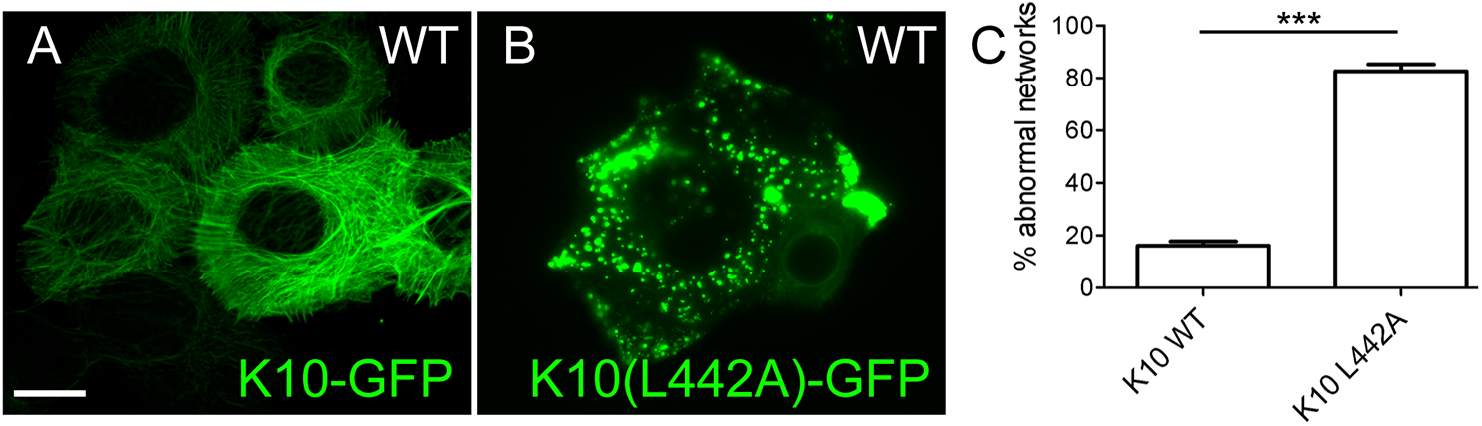
WT keratinocytes were tranfected with either WT keratin 10-GFP or with a mutant that cannot interact with Ndel1, keratin 10-L442A-GFP. Quantitation of cells with abnormal networks is shown in (D). Error bars represent SEM.

**Figure 3, Supplement 1.**
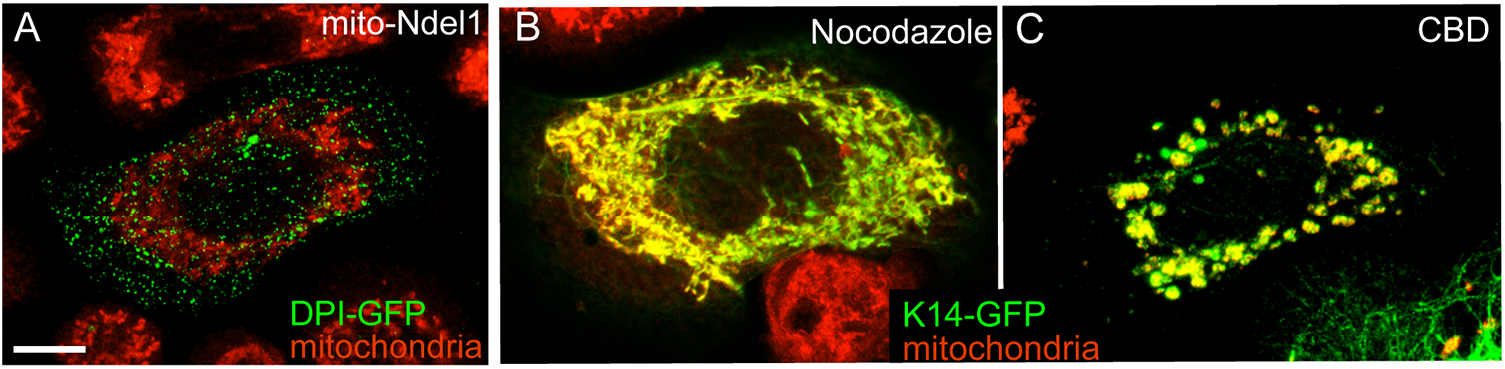
(A) WT keratinocytes were transfected with DPI-GFP (green) and mito-Ndel1. Cells were treated with MitoTracker-Red to label mitochondria. (B,C) WT keratinocytes transfected with mito-Ndel1 and K14-GFP were treated with either Nocodazole (B), to disrupt microtubules, or cilibrevin D (CBD, to inhibit dynein). Scale bar is 10 μm.

**Figure 3, Supplement 2.**
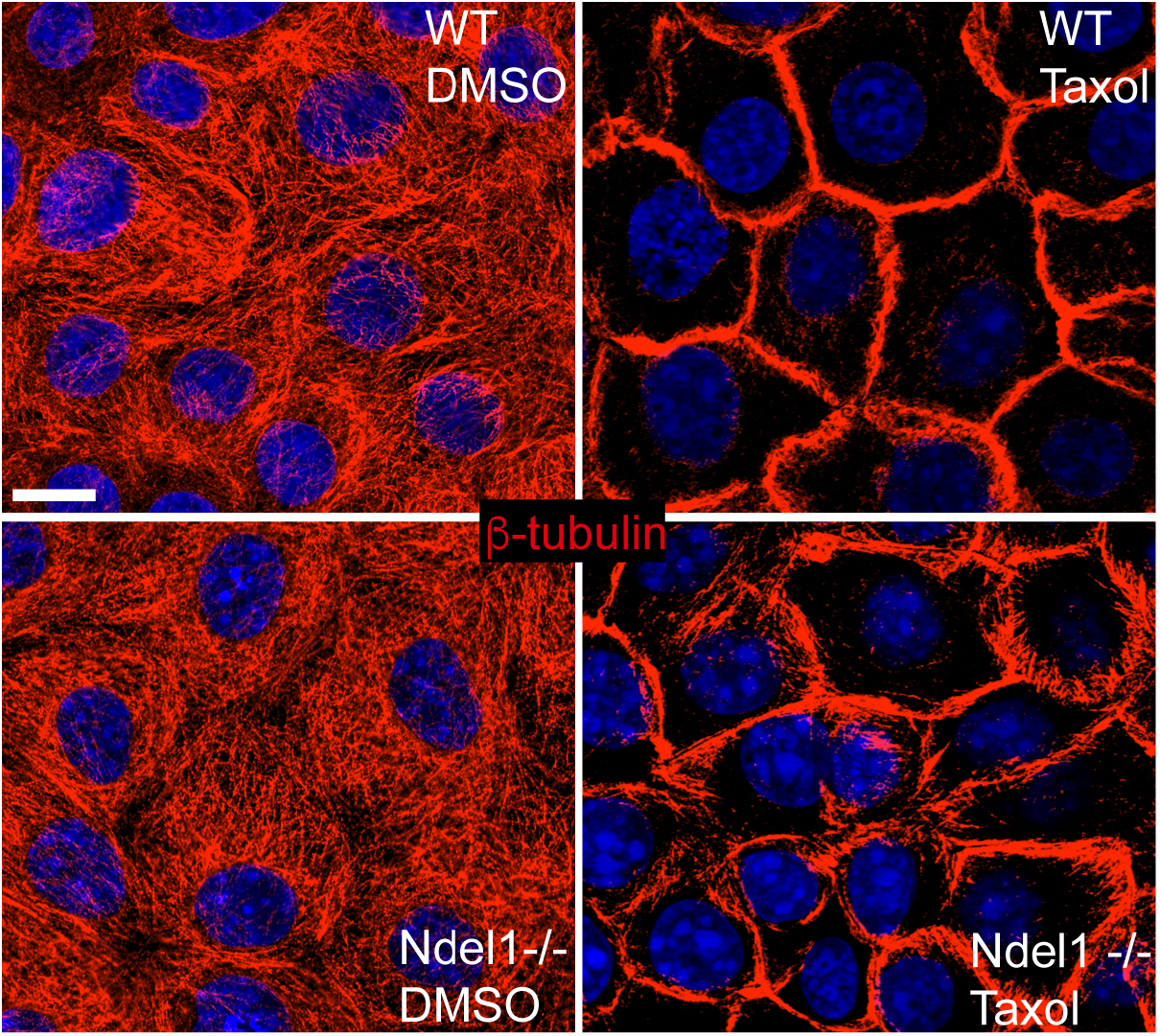
Microtubule reorganization to the cell cortex is normal in Ndel1-/-cells. Taxol was used to stabilize microtubules, which promotes their cortical organization in keratinocytes.

**Figure 4, Supplement 1.**
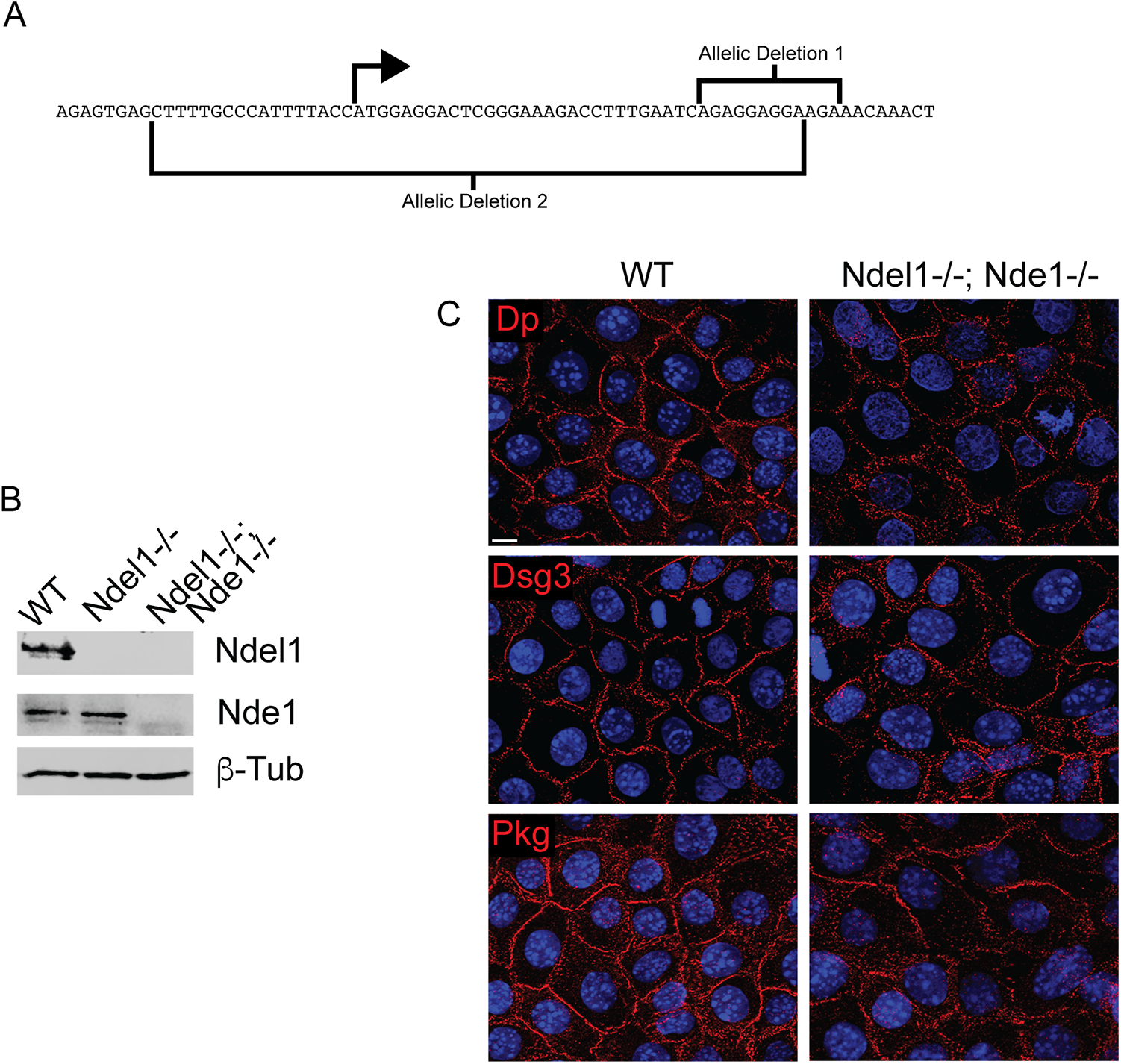
Generation of Nde1-/-cells. (A) Diagram of the region targeted by CRISPR and the two independent mutations generated in the clonal line used. (B) Western blot of lysates from control, Ndel1 null and Ndel1;Nde1 double null cell. β-tubulin is used as a loading control. (C) Immunofluorescence of desmosomal proteins, as indicated, in control and Nde1;Nde1 double null cells. Scale bar is 10 μm.

